# POAnoise: A Graph-based Denoising Pipeline for Amplicon Sequencing Data

**DOI:** 10.64898/2026.09.14.751445

**Authors:** Muhammad Ardiyansyah, Brendan Furneaux, Otso Ovaskainen

**Author notes:** **Corresponding author** **(MA)**.

## Abstract

High-throughput DNA metabarcoding enables large-scale biodiversity assessment by identifying taxa from environmental samples, but its accuracy critically depends on denoising methods that separate true biological variation from PCR and sequencing errors. A persistent challenge is robust reconstruction of sequence diversity across abundance distributions, where low-abundance variants are particularly difficult to recover.

We introduce **POAnoise**, a graph-based denoising framework that uses Partial Order Alignment (POA) to model relationships among noisy sequencing reads. POAnoise incrementally constructs sequence graphs that represent substitutions and indels, and derives consensus sequences from graph-supported paths using a weighted consensus strategy. By combining graph-based alignment with abundance-aware clustering, the method provides a structured way to reconstruct sequence variants from noisy amplicon data across heterogeneous abundance regimes.

We evaluated POAnoise on simulated ITS and 16S datasets and compared its performance with established denoising methods, DADA2 and UNOISE3, across multiple parameter settings. Across the benchmark datasets, POAnoise generally achieved higher F_1_-scores and exhibited more stable performance across parameter configurations. In abundance-aware analyses, POAnoise showed reconstruction ratios closer to unity and reduced abundance-dependent deviation compared with DADA2, while remaining broadly comparable to UNOISE3 across most abundance classes. Overall, these results indicate that POAnoise can provide a robust alternative for amplicon denoising.

**Author Summary:** DNA sequencing allows us to study the diversity of microorganisms in environmental samples, but errors introduced during laboratory and sequencing processes can make it difficult to distinguish real biological sequences from noise. This is particularly challenging when some sequences are much more abundant than others. In this study, we developed POAnoise, a graph-based approach for reducing errors in amplicon sequencing data by using relationships among similar sequences to reconstruct the underlying biological diversity. We tested POAnoise on simulated bacterial and fungal sequencing data and compared it with commonly used denoising methods. We found that POAnoise generally recovered true sequences accurately and showed consistent performance across different settings and abundance levels. Our results suggest that considering relationships among sequences can improve the reliability of amplicon sequencing analyses and may help researchers obtain more consistent estimates of microbial diversity.

## 1 Introduction

High-throughput DNA sequencing (HTS) has enabled large-scale characterization of genetic variation and microbial diversity. However, sequencing errors can obscure the distinction between true biological sequence variants and technical artifacts, particularly in amplicon sequencing. Illumina sequencing is dominated by nucleotide substitutions, although insertions and deletions can also occur [15]. Accurate denoising is therefore essential for recovering biological sequence variants and for avoiding downstream errors in ecological and diversity analyses [7].

Before the advent of modern denoising methods, sequencing reads were commonly clustered based on a fixed sequence-similarity threshold [8, 20]. Such threshold-based clustering can merge distinct biological sequence variants with sequencing artifacts, limiting resolution at the single-nucleotide level [4]. This limitation prompted the development of more advanced denoising tools such as the Divisive Amplicon Denoising Algorithm (DADA) and its improved version, DADA2, which model sequencing errors to infer exact sequence variants (ESVs or ASVs) [5, 19]. In contrast with fixed-threshold clustering approaches, an obvious advantage of using these ASV approaches is the consistency of ASVs as sequence labels that can be used across studies without the need for re-clustering [3]. These ASV approaches rely on different statistical models to distinguish true biological sequences from the spurious ones.

DADA2 represents a model-based approach in which an empirical error model is learned from the sequencing data and used to infer ASVs with single-nucleotide resolution [5]. Such error-model-based methods provide a principled framework for distinguishing sequencing errors from biological variation, but their performance depends on the adequacy of the assumed error model and on parameter choices. In particular, variants whose abundance or sequence differences are difficult to distinguish from the estimated error process may be challenging to recover.

Other denoising approaches use different signals to distinguish biological variants from sequencing errors. UNOISE3, for example, relies primarily on sequence abundance and similarity heuristics to identify denoised sequence variants [9]. Deblur instead uses a pre-trained error model for rapid and standardized processing of short-read amplicon data [2]. Thus, existing denoising methods exploit statistical error profiles, sequence similarity, abundance, or combinations of these signals to identify putative biological variants.

Existing denoising methods also differ in how they represent relationships among sequences. DADA2 and UNOISE rely primarily on pairwise sequence comparisons to identify relationships between candidate variants and potential sequencing errors, whereas Deblur does not explicitly use sequence alignment as part of its denoising framework. Multiple sequence alignment provides a different representation by capturing shared sequence structure across a collection of reads simultaneously. In particular, sequences originating from the same biological variant may follow similar paths through a multiple alignment, while sequencing errors can appear as localized deviations from these shared patterns. This raises the question of whether multiple sequence alignment can provide a useful structural signal for distinguishing biological sequence variants from sequencing errors.

Partial Order Alignment (POA) provides a natural framework for representing such alignment structure. POA represents a multiple sequence alignment as a directed acyclic graph (DAG), in which nodes represent nucleotide positions and edges encode allowable transitions between positions [13]. This graph representation naturally captures alternative alignment paths and branching sequence variation, including substitutions and indels, while preserving the relationships among sequences in the alignment. POA has been particularly useful for consensus generation and long-read sequence analysis, but its use as a graph-based representation for denoising short-read amplicon data has received comparatively less attention [13, 18, 22].

In this study, we ask whether the structural information encoded by a partial-order alignment graph can provide a useful signal for distinguishing true biological sequence variants from sequencing errors. To address this question, we introduce **POAnoise**, a graph-based denoising framework that uses POA graphs to represent sequence relationships and abundance weighted edge-traversal patterns to characterize sequence similarity. These graph-based features are used to cluster noisy sequences into groups with similar alignment structure, after which a separate POA is constructed for each cluster and used to generate a consensus sequence. This framework therefore combines alignment structure, observed sequence abundance, and graph-based consensus reconstruction in a single denoising pipeline.

We evaluate POAnoise on simulated ITS and 16S amplicon datasets with controlled sequencing errors and abundance variation, and compare its performance with DADA2 and UNOISE3 across multiple parameter configurations. We specifically assess sequence-level precision, recall, F_1_-score, error rates, and abundance-aware reconstruction to determine whether the graph-based representation provides a robust signal for denoising under different sequence and abundance regimes.

## 2 Methods

### 2.1 POAnoise Pipeline

*Partial Order Alignment* (POA) is a graph-based framework for representing and aligning multiple biological sequences in a flexible and memory-efficient manner. Originally introduced by Lee et al. (2002) [13], POA encodes a multiple sequence alignment as a *partial order graph* (POG), where nodes correspond to nucleotides (or amino acids) and edges represent observed alignment transitions between bases. In contrast to the conventional matrix representation of a multiple sequence alignment, the graph representation makes alternative alignment paths and their relationships explicit. This allows sequence variation, including substitutions and insertions or deletions, to be represented as branching paths within the same alignment structure. Building upon this foundation, we introduce **POAnoise** which is a graph-based denoising method that incrementally constructs a POA graph from noisy DNA sequences. Figure 1 visualizes the POAnoise pipeline. In POAnoise, as in other denoising workflows, the input noisy sequences are first filtered based upon their quality scores and then are dereplicated to obtain unique representation of the noisy data. Afterwards, POA graphs of the dereplicated and filtered sequences are constructed. From these graphs, we can then obtain a weighted feature matrix that encodes the sequence and POA edges co-occurrence patterns. This feature matrix are translated to a normalized pairwise distance matrix which is used to cluster our noisy sequences based on feature similarities. From each cluster, a set of consensus sequences representing the cluster is computed. The collection of all consensus sequences along with the number of reads supporting each one is the output of our denoising algorithm. The source code, benchmarking scripts, and analysis pipeline for POAnoise are publicly available at: https://github.com/ardiyansyah13/POAnoise-implementation

**Figure 1.**
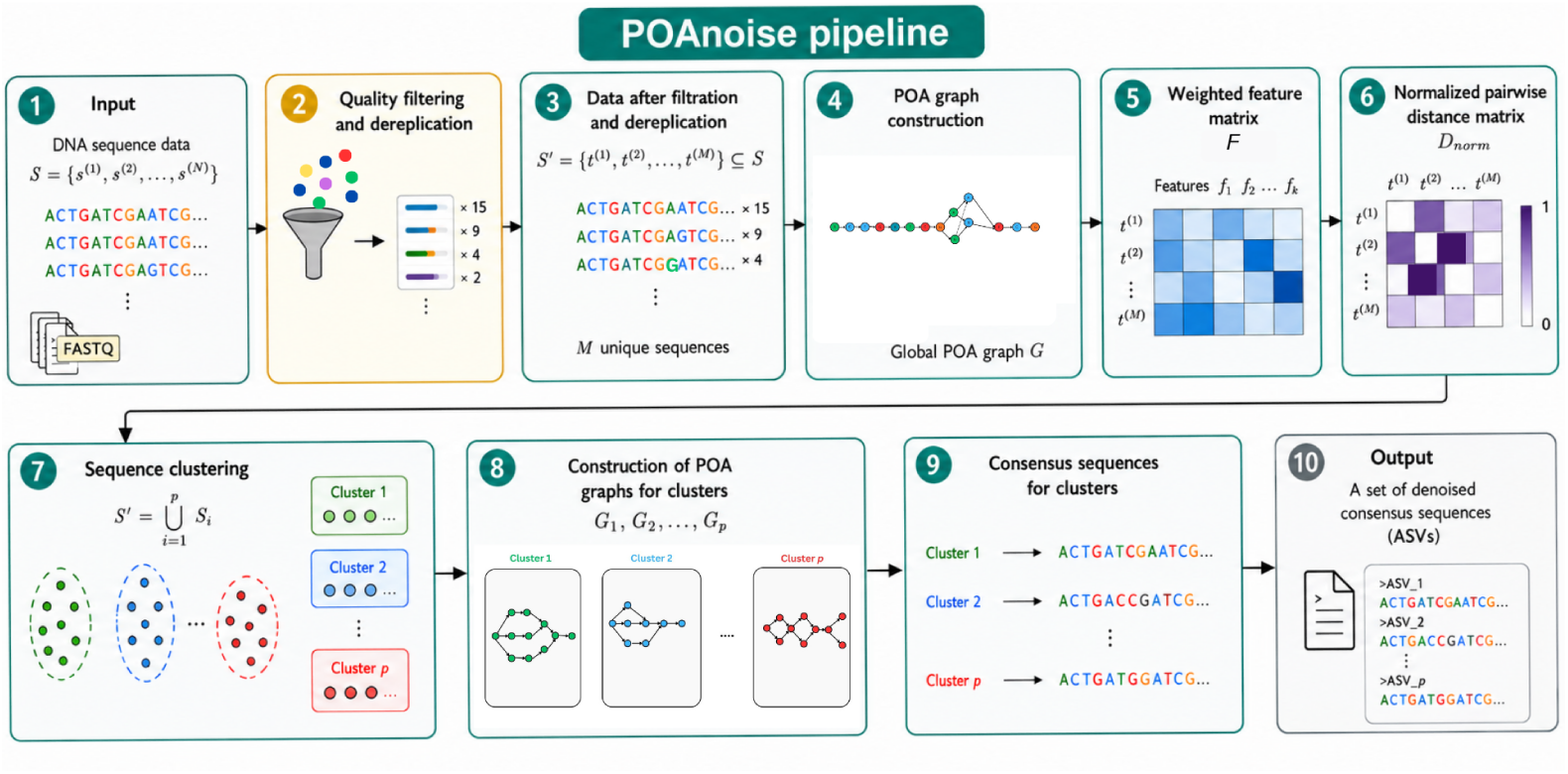
Overview of the POAnoise pipeline. (1) Input DNA sequencing reads are provided as a set of raw sequences. (2) The reads undergo quality filtering and dereplication, retaining high-quality unique sequences together with their abundances. (3) The resulting set of *M* unique sequences is used as input for the subsequent analysis. (4) A global partial order alignment (POA) graph *G* is constructed from the unique sequences. (5) Sequence features are extracted from the POA graph to construct a weighted feature matrix *F*. (6) The weighted feature matrix is used to compute a normalized pairwise distance matrix *D*_norm_ between sequences. (7) The unique sequences are clustered into *p* groups based on the normalized pairwise distances. (8) A separate POA graph *G*_1_*, …, G_p_* is constructed for each sequence cluster. (9) A consensus sequence is generated from each cluster-specific POA graph. (10) The resulting consensus sequences constitute the final set of denoised amplicon sequence variants (ASVs).

The repository includes implementations of the POA-based denoising framework, parameter sweep experiments, statistical analyses, and figure-generation scripts used in this study. All computations and analyses in this study were performed on a laptop workstation equipped with an Intel(R) Core(TM) Ultra 7 165U processor featuring 12 cores and 14 logical CPUs, with a maximum clock speed of 4.9 GHz, running a 64-bit Linux operating system on x86 64 architecture.

#### 2.1.1 Partial Order Alignment (POA)

Formally, let *S* = *{s*^(1)^*, s*^(2)^*, …, s*^(^*^N^*^)^*}* be a set of input noisy DNA sequences. The POA graph *G* = (*V, E*) associated to *S* is a directed acyclic graph (DAG) where each node *v ∈ V* represents a nucleotide base of a sequence in *S*, and each directed edge *e ∈ E* represents an allowable transition between bases. Each node also stores the set of sequences that traverse it. The POA *G* is initialized from the first sequence *s*^(1)^ as a linear chain of nodes, and for each subsequent sequence *s*^(^*^i^*^)^, the *dual-affine Needleman–Wunsch (NW)* algorithm [10, 16] is applied to align it to the existing graph and update *G*. The alignment routine employs dynamic programming (DP) with multiple gap penalty states. The nodes of *G* are first topologically sorted into distinct positions, and the DP states represent alignment ending in a base match or mismatch, insertion with the first gap model (*I*_1_), insertion with the second gap model (*I*_2_), deletion with the first gap model (*D*_1_), or deletion with the second gap model (*D*_2_). The nucleotide scoring function assigns a score of 1 to a match and *−*1 to a mismatch. Gap penalties are represented by the upper envelope of two affine gap-score functions,

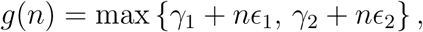

where *n* denotes the gap length, (*γ*_1_*, ɛ*_1_) defines the first gap regime, and (*γ*_2_*, ɛ*_2_) defines the second gap regime. The use of the maximum is consistent with score maximization: for a given gap length, the affine component pro-viding the larger alignment score is selected. This two-regime construction is designed to accommodate different gap-length regimes within the same alignment model. After the DP matrices have been computed, traceback identifies the optimal alignment path between the input sequence and the POA graph, and the POA is updated according to the resulting alignment.

#### 2.1.2 POA-Based Sequence Feature Extraction and Clustering

Once all unique sequences have been incorporated into the global POA graph, we represent each sequence by its pattern of edge traversals through the graph. This representation provides a graph-based description of sequence similarity that is sensitive not only to nucleotide composition but also to the structural paths taken through the alignment graph.

Let *S* = *{s*^(1)^*, …, s*^(^*^M^*^)^*}* denote the set of *M* dereplicated sequences, and let *n_i_* denote the observed abundance of sequence *s*^(^*^i^*^)^. We define the weighted feature matrix 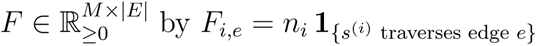, where **1***_{·}_* is the indicator function. Thus, each row of *F* represents one unique sequence, while each column represents an edge of the global POA graph.

The key assumption underlying this representation is that sequencing errors and biological variation may both introduce substitutions or indels, but their patterns are expected to differ across reads. Sequencing errors are typically introduced independently during read generation, so the specific substitutions or indels associated with errors are not expected to be consistently shared across many reads. In contrast, a true biological variant is expected to generate multiple reads carrying the same sequence differences. Consequently, reads originating from the same biological variant are expected to follow similar paths through the POA graph, whereas error-containing sequences may deviate from these paths in different locations.

The edge-traversal representation captures this distinction by encoding the alignment path followed by each sequence. Sequences sharing a biological variant therefore tend to have similar patterns of edge traversal, while sequences carrying independent sequencing errors tend to produce more localized and heterogeneous deviations. The resulting feature vectors provide a representation of sequence relationships based on their shared multiple-alignment structure rather than solely on direct pairwise sequence similarity. Moreover, once the global POA graph has been constructed, each sequence can be represented by its traversal of this common graph. Thus, for *M* sequences, constructing the feature representation requires *O*(*M*) sequence–graph alignments, rather than *O*(*M* ^2^) pairwise sequence–sequence alignments required for an all-pairs comparison.

Abundance is incorporated into the representation because repeated observations provide additional support for a sequence traversal pattern. A high-abundance sequence therefore contributes more strongly to the corresponding graph features than a low-abundance sequence. The resulting representation combines alignment structure with observed abundance and is used for clustering sequences according to their shared POA traversal patterns.

For two sequences *s*^(^*^i^*^)^ and *s*^(^*^j^*^)^, similarity between their feature vectors is quantified using cosine similarity,

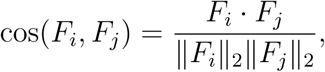

and the corresponding cosine distance is *D_ij_* = 1 *−* cos(*F_i_, F_j_*). Because the entries of *F* are non-negative, *D_ij_* lies in [0, 1]. We therefore use the resulting distance matrix directly for subsequent clustering.

We then apply agglomerative hierarchical clustering with single linkage to the distance matrix *D*. Single linkage is particularly suitable for the POA representation because sequences belonging to the same underlying variant can form connected groups through intermediate error-containing sequences. Thus, clustering based on the minimum inter-sequence distance can retain connected sequence neighborhoods rather than requiring every pair of sequences within a cluster to satisfy the same distance criterion.

Clusters are obtained by cutting the resulting dendrogram at a threshold *t ∈* [0, 1]. In the benchmarking experiments, a range of values of *t* is considered in order to evaluate the sensitivity of POAnoise to the clustering parameter. For a fixed threshold *t*, two sequences are connected whenever their hierarchical clustering distance is at most *t*, and the resulting connected groups define the sequence clusters *S* = *{S*_1_*, …, S_p_}, S_k_ ⊆ S*.

The purpose of this clustering step is to partition the global POA into groups of sequences with similar graph-traversal structure. Under the denoising model, sequencing-error variants are expected to occur as low-abundance sequences located near more strongly supported sequence patterns, whereas distinct biological variants are expected to produce separate traversal patterns. The clustering step therefore provides the candidate sequence groups within which consensus reconstruction is performed.

#### 2.1.3 Consensus Construction for Clusters

For each cluster *S_k_* (1 *≤ k ≤ p*), a cluster-specific POA *G_k_* is constructed by realigning the sequences in *S_k_* using the same dual-affine Needleman– Wunsch alignment described above. We construct a new POA rather than simply subsetting the global graph *G*, because the global graph contains alignment paths supported by sequences from other clusters. Re-alignment therefore allows *G_k_* to represent the sequence variation and edge-traversal support specific to cluster *S_k_*. Each edge (*u, v*) *∈ E*(*G_k_*) is assigned a weight *w*(*u, v*) corresponding to the number of sequences in *S_k_* that traverse that edge.

Let *C* = (*v*_1_*, …, v_m_*) denote an admissible source-to-sink path in *G_k_*. Following the heaviest-bundle consensus procedure introduced for POA [13], the support of a path is defined as

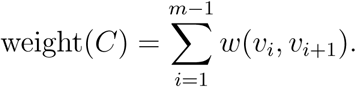

The consensus sequence is obtained from the path

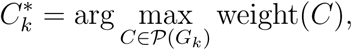

where *P*(*G_k_*) denotes the set of admissible paths through *G_k_*. Thus, the selected path corresponds to the sequence variant with the largest cumulative traversal support within the cluster.

If multiple paths attain the same maximum weight, they are retained as candidate consensus sequences. Finally, identical consensus sequences obtained from different clusters are merged to produce the final set of denoised sequence variants.

### 2.2 Evaluating the Performance of POAnoise

To systematically assess the performance of POAnoise, we designed a controlled evaluation pipeline that begins with curated reference sequences and proceeds through noise simulation, denoising, and quantitative comparison against ground truth. The workflow integrates realistic sequencing error models, abundance variation, and taxonomic diversity to ensure that benchmarking reflects practical use cases. An overview of this evaluation procedure is illustrated in Figure 2, which summarizes the key stages from dataset construction to performance assessment.

**Figure 2.**
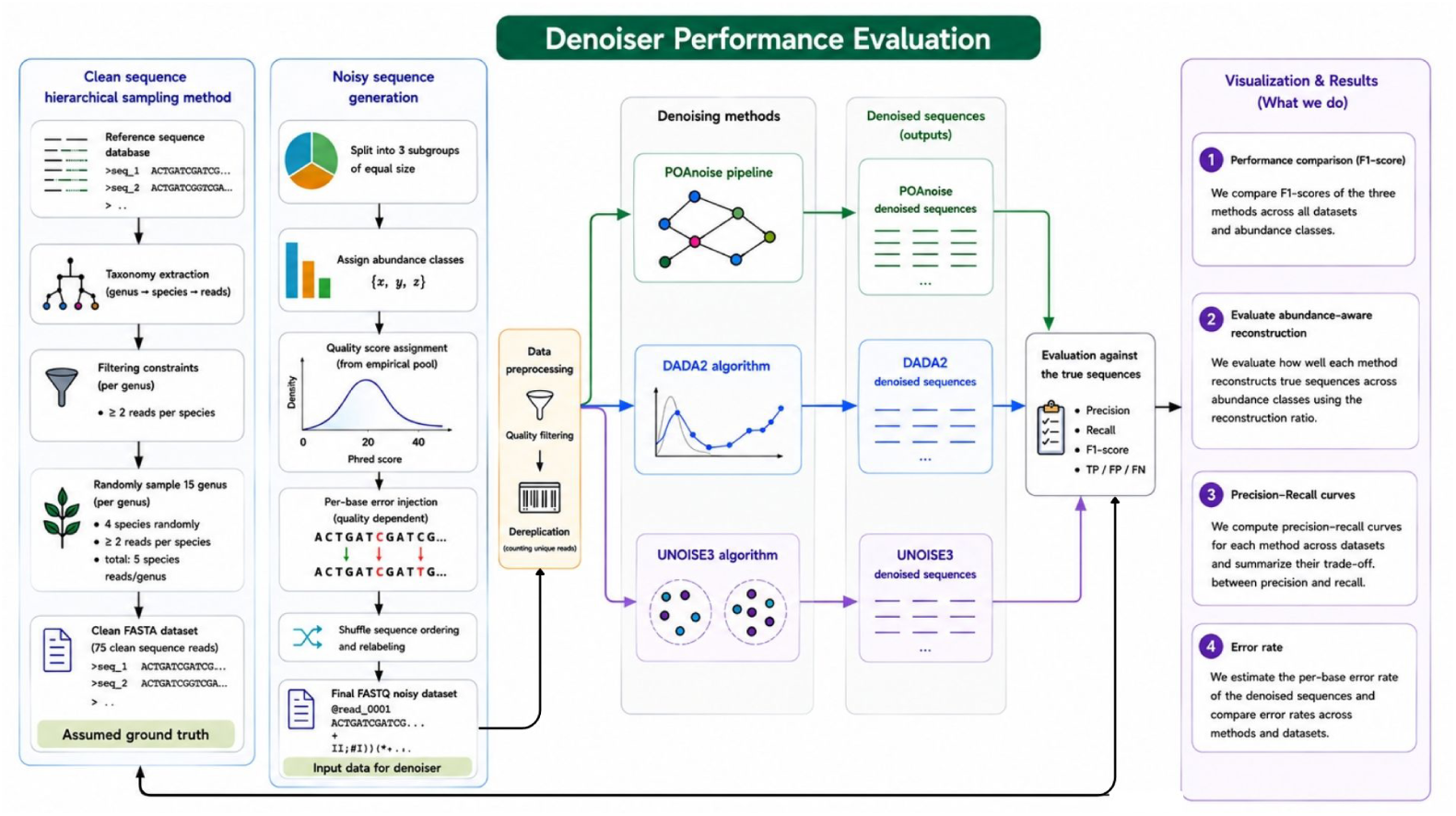
Overview of the denoiser performance evaluation workflow. Clean sequences are used to generate quality-dependent noisy datasets with controlled abundance classes, which are processed by POAnoise, DADA2, and UNOISE3. The resulting denoised sequences are evaluated against the true sequences using precision, recall, F1-score, reconstruction ratio, and error rate.

In the evaluation, DADA2 and UNOISE3 were selected because they are widely used denoising methods for amplicon sequencing data and represent distinct denoising strategies. DADA2 uses an explicit statistical error model, whereas UNOISE3 relies primarily on abundance- and similarity-based heuristics. Comparing POAnoise with these two methods therefore provides a benchmark against established approaches representing complementary denoising paradigms.

#### 2.2.1 Case Study with ITS Data

The first benchmark dataset used to evaluate POAnoise was constructed from the *SH General Release Dataset* of the UNITE database [1]. We selected ITS sequences containing the ITS3 primer (GCATCGATGAAGAACGCAGC) and trimmed each sequence to start at the primer site, retaining the full sequence downstream of the primer. To ensure taxonomic diversity, we retained species with at least two sequences and genera containing at least four such species. From the resulting set, 15 genera were randomly selected. For each genus, four species were sampled: one species contributed two distinct sequences, while the remaining three species contributed one sequence each, resulting in five reference sequences per genus and 75 reference sequences in total. Duplicate nucleotide sequences were removed before sampling to ensure that the two sequences selected from the same species were non-identical.

#### 2.2.2 Case Study with 16S Data

The second benchmark dataset was constructed from the SILVA SSURef NR99 database (Release 138; https://ftp.arb-silva.de) [17, 23], which provides full-length 16S rRNA sequences with standardized taxonomic annotations. The V4 hypervariable region was extracted using the 515F-Y primer (GTGYCAGCMGCCGCGGTAA) in an IUPAC-aware manner, retaining sequences with valid primer matches. For each retained sequence, a 250 bp fragment downstream of the primer site was extracted. To obtain a taxonomically diverse reference set comparable to the ITS benchmark, we selected genera containing at least four distinct species, with each species represented by at least two sequences.

#### 2.2.3 Common Simulation and Abundance Design

The same simulation framework was applied to both the ITS and 16S reference datasets. This common design was used to ensure that differences observed between the two marker types were not introduced by differences in the sequencing-error or abundance-generation procedures.

To incorporate realistic sequencing-quality variation, Phred quality scores were derived from an empirical Illumina FASTQ dataset. The resulting empirical Phred quality-score profiles were independently assigned to simulated reads. For each read, the sequence and its assigned quality profile were truncated to the shorter of the two lengths, ensuring that every nucleotide in the simulated read was associated with a corresponding quality score. This procedure preserves the empirical distribution of quality scores while allowing the simulated reads to vary in length according to the available sequence and quality-profile lengths.

Noisy reads were generated using a Phred score–dependent substitution error model. For each nucleotide position with quality score *q*, the sequencing error probability was defined as

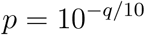

following the nominal definition of the Phred quality score [6]. With probability *p*, the original nucleotide was replaced uniformly at random by one of the three alternative nucleotides in *{A, C, G, T} \* true base. Errors were introduced independently across nucleotide positions, conditional on the assigned empirical quality profile.

To evaluate denoising performance under heterogeneous sequence abundances, the reference sequences in each marker dataset were partitioned into three equal-sized groups. Each group was assigned a fixed abundance level, corresponding to low-, intermediate-, and high-abundance sequences. For the ITS benchmark, the three abundance regimes were ITS_A:*{*1, 4, 16*}*, ITS_B:*{*4, 16, 64*}*, and ITS_C:*{*16, 64, 256*}*. The corresponding 16S datasets were generated using the same three-level abundance design, yielding the datasets 16S_A, 16S_B, and 16S_C. Within each dataset, reference sequences were replicated according to their assigned abundance level. Each replicate was generated independently, including independent sampling of a quality profile and independent introduction of sequencing errors. After simulation, all reads were randomly shuffled to remove ordering effects introduced during dataset generation.

Finally, each simulated read was assigned a unique identifier, and metadata linking the read to its originating reference sequence and assigned abundance class were stored alongside the resulting FASTQ files. The resulting datasets therefore provide known ground truth at both the sequence and abundance levels, enabling direct evaluation of sequence recovery and abundance-aware reconstruction.

#### 2.2.4 Applying POAnoise, DADA2, and UNOISE3 to the Case Studies

To provide a fair and controlled comparison between POAnoise and established denoising algorithms, both DADA2 and UNOISE3 were applied to the same synthetic noisy datasets used in each case study. All three algorithms implemented with varying certain parameters were therefore applied to the same noisy input datasets to systematically exploring the impact of parameter choices on denoising performance. The resulting denoised sequences were then used for precision and recall computation at the sequence variant levels described in the following sections.

Raw FASTQ files containing noisy datasets were initially filtered using a maximum expected-error threshold of maxEE = 2, discarding reads with excessive predicted sequencing error. This is a commonly used default setting in DADA2-based amplicon workflows [5]. The remaining reads were dereplicated, collapsing identical sequences while retaining their observed abundances. These unique sequences and their abundances were then used as input to the POAnoise pipeline.

For all POAnoise analyses, the dual-affine Needleman–Wunsch alignment used the following fixed scoring scheme: match = 1, mismatch = *−*1, with the two affine gap components parameterized as (*γ*_1_*, ɛ*_1_) = (*−*1*, −*1), (*γ*_2_*, ɛ*_2_) = (*−*5, 0). Thus, the resulting gap score is

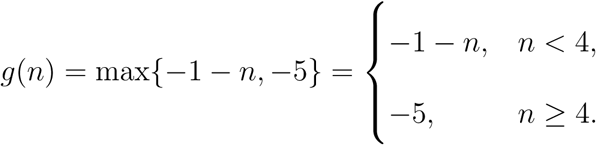

Thus, short gaps incur an increasingly severe penalty with increasing length, whereas gaps of length four or greater are assigned the constant score *−*5.

The clustering threshold was the primary POAnoise parameter varied in the benchmark experiments. We considered thresholds *t ∈ {*0.01, 0.06, 0.11*, …* covering a broad range of pairwise-distance cutoffs. For each value of *t*, the sequences were clustered according to the normalized POA-based distance matrix, and a separate consensus sequence was reconstructed for each resulting cluster. This parameter sweep was used to evaluate the sensitivity of POAnoise performance to the clustering threshold.

DADA2 was run using its standard pipeline for single-end reads. The DADA2 pipeline consisted of the following steps: (1) quality filtering and trimming, (2) dereplication, (3) error model learning, and (4) DADA2 core sample inference. Quality filtering and trimming are performed with key parameters varied across runs: maxEE = 2, truncQ = 2, read truncation to 300 bp for ITS and 250 bp for 16S and adjusted nbases = 10^7^ to control the number of bases processed. DADA2 error-model learning and sequence inference were then performed together using the core DADA2 procedure. We varied the OMEGA A parameter over *{*10*^−^*^50^, 10*^−^*^40^, 10*^−^*^30^*}* and the BAND SIZE parameter over *{*16, 24, 32*},* to assess the sensitivity of DADA2 inference to these settings.

UNOISE3 was executed using USEARCH (v11). For each noisy dataset, multiple parameter combinations were explored to evaluate performance. Following the dereplication stage of noisy reads using -fastx uniques with -sizeout to retain abundance information and -relabel to standardize sequence identifiers, the core denoising stage is performed via the -unoise3 command to generate ZOTUs (zero-radius OTUs), representing denoised sequence variants. The -minsize parameter controlling the minimum abundance of sequences considered was varied *∈ {*1, 2, 4, 6, 8*}*, and the -abskew parameter controlling maximum allowed abundance skew between sequences was varied *∈ {*2, 3, 4, 5*}*.

#### 2.2.5 Performance Evaluation

Sequence-level performance was evaluated by comparing the denoised sequences with a curated reference sequence set representing the known ground truth for each synthetic dataset. Let *R* = *{r*_1_*, …, r_K_}* denote the set of *K* true reference sequences and *A* = *{a*_1_*, …, a_L_}* the set of *L* denoised sequences produced by a given method and parameter configuration. For each denoised sequence *a_i_*, we computed its global pairwise alignment against every reference sequence *r_j_* using the Needleman–Wunsch global alignment implemented in Biopython. Let *s_ij_ ∈* [0, 1] denote the sequence similarity between *a_i_* and *r_j_*, defined as the proportion of identical residues in the aligned region after excluding terminal overhangs. The best reference match for *a_i_* was defined as

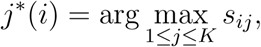

with corresponding maximum similarity *s_i_* = *s_i,j_∗*_(_*_i_*_)_.

A denoised sequence was considered a valid reconstruction if *s^∗^ ≥ τ,* where *τ* denotes the predefined similarity threshold. Denoised sequences with *s^∗^ < τ* were classified as false positives. The threshold *τ* was fixed throughout the benchmark. Because multiple denoised sequences can potentially match the same reference sequence, reference-level recovery was evaluated separately. A reference sequence *r_j_* was considered recovered if at least one denoised sequence satisfied *s_ij_ ≥ τ.* The number of recovered reference sequences was therefore

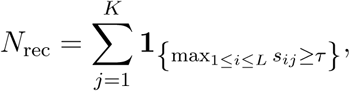

and the number of unrecovered reference sequences was *FN* = *K − N*_rec_.

Precision was calculated at the denoised-sequence level as Precision = 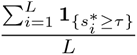. Recall was calculated at the reference-sequence level as Recall = 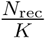 . The corresponding F_1_-score was calculated as the harmonic mean of the precision and the recall score. For the primary performance evaluation, we used a predefined similarity threshold of *τ* = 0.99, such that a denoised sequence was considered a valid reconstruction when its similarity to the best-matching reference sequence was at least 99%. Additionally, we evaluated sequence reconstruction accuracy using error rates calculated as the proportion of mismatched positions after global alignment between each denoised sequence and its closest reference sequence.

To ensure a comprehensive assessment across parameter space, multiple denoised outputs were generated for each dataset and method. POAnoise was evaluated using 20 parameter configurations corresponding to combinations of clustering thresholds. DADA2 was evaluated using 9 parameter configurations spanning the OMEGA_A and BAND_SIZE parameters. UNOISE3 was evaluated using 20 parameter configurations defined by combinations of the -minsize and -abskew parameters.

To statistically assess differences in performance distributions across methods, F_1_-scores were analyzed using a nonparametric framework due to significant deviations from normality as indicated by the Shapiro–Wilk test [7, 21]. Overall differences among denoisers within each dataset were evaluated using the Kruskal–Wallis test [12], a rank-based nonparametric analogue of one-way ANOVA. When significant differences were detected, pairwise comparisons were performed using the Mann–Whitney U test [14] with Holm correction for multiple hypothesis testing [11].. A significance level of 0.05 was applied to corrected p-values.

For the abundance-aware evaluation, reconstructed sequences were additionally analyzed according to the abundance of their originating reference sequences. As described in Section 2.2, reference sequences were partitioned into predefined abundance classes representing high-, medium-, and low-abundance taxa. For each denoiser, similarity distributions of reconstructed sequences were compared within each abundance class to assess abundance-dependent performance variation.

## 3 Results

Across all datasets, POAnoise consistently achieved the highest median and mean F_1_-scores relative to UNOISE3 and DADA2 (Figure 3a). POAnoise also showed relatively compact distributions across parameter configurations, indicating stable performance across the tested settings. The F_1_-score trends across individual parameter configurations further illustrate this consistency, with POAnoise maintaining high performance across datasets, whereas UNOISE3 showed greater sensitivity to parameter choice (Figure 3b). DADA2 generally yielded the lowest F_1_-scores, particularly for the ITS datasets. Pairwise comparisons supported these differences for most method pairs; the main exception was DADA2 versus UNOISE3 for 16S A, for which the difference was not significant.

**Figure 3.**
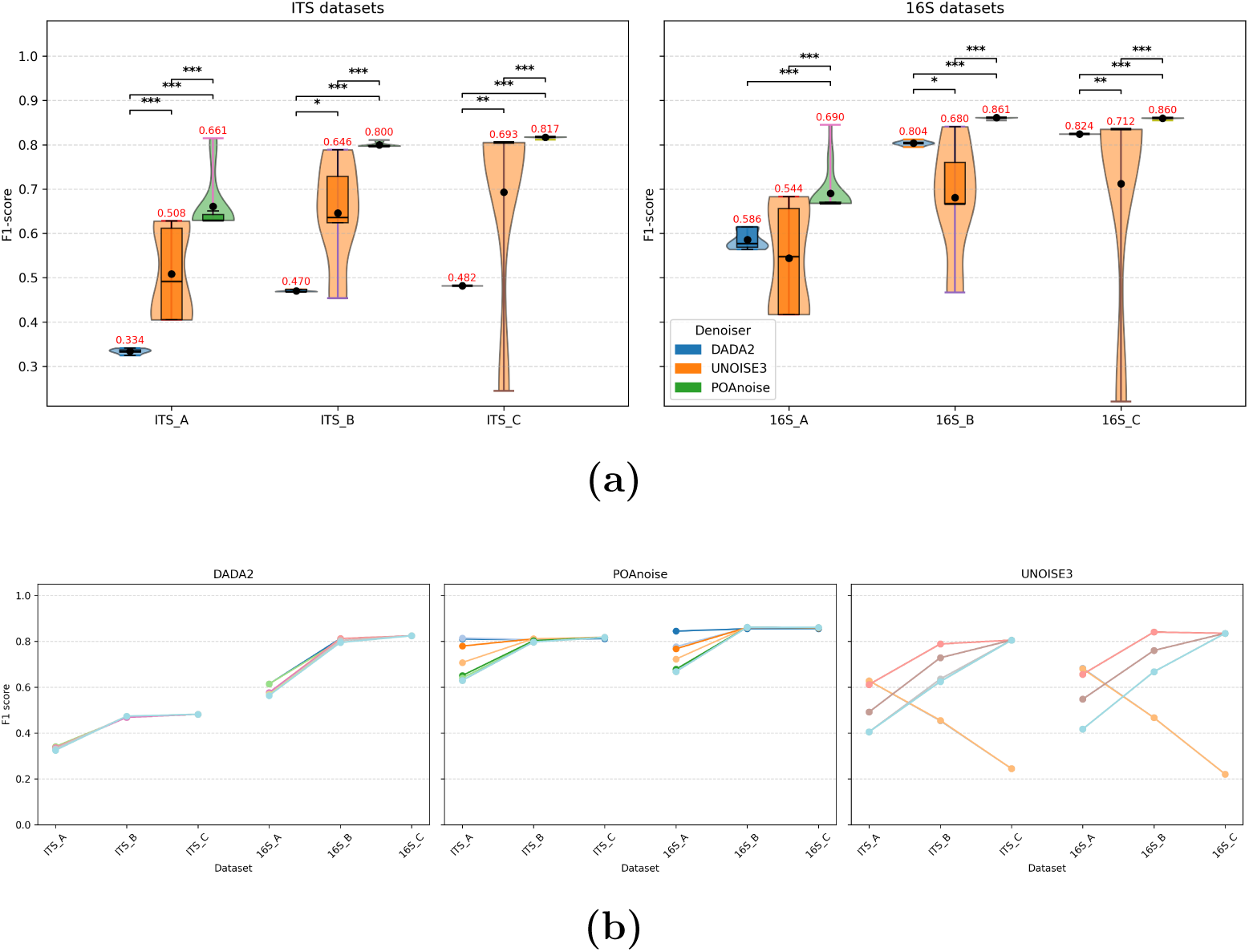
Performance comparison of denoising methods across ITS and 16S datasets under varying parameter configurations. (a) Distribution of F_1_-scores across denoising methods and parameter settings. Each violin shows the distribution of F_1_-scores obtained across parameter configurations for a given denoising method and dataset. The embedded boxplot shows the median, interquartile range (IQR), and whiskers extending to the most extreme values within 1.5*×*IQR. Black dots indicate the mean F_1_-score, with the corresponding mean value shown in red above each distribution. Horizontal brackets indicate significant pairwise differences between denoising methods based on Holm-adjusted Mann–Whitney tests. One, two, and three stars indicate adjusted *p <* 0.05, *p <* 0.01, and *p <* 0.001, respectively; nonsignificant comparisons are not annotated. (b) F_1_-score trends for DADA2, UNOISE3, and POAnoise across ITS and 16S datasets. Lines indicate the F_1_-score obtained for individual parameter configurations across datasets, with colors corresponding to parameter configurations.

The precision–recall analysis showed a similar pattern (Figure 4). POAnoise configurations consistently clustered toward the upper-right region of the precision–recall space, corresponding to simultaneously high precision and recall. Consequently, POAnoise points generally occupied higher F_1_-score contours than those of UNOISE3 and DADA2, indicating a more favorable balance between the two measures. DADA2 configurations were generally concentrated toward lower recall, particularly for the ITS datasets, whereas UNOISE3 showed a wider spread across the precision–recall space. These results are consistent with the overall F_1_-score distributions and indicate that POAnoise maintains strong performance across the tested parameter configurations.

**Figure 4.**
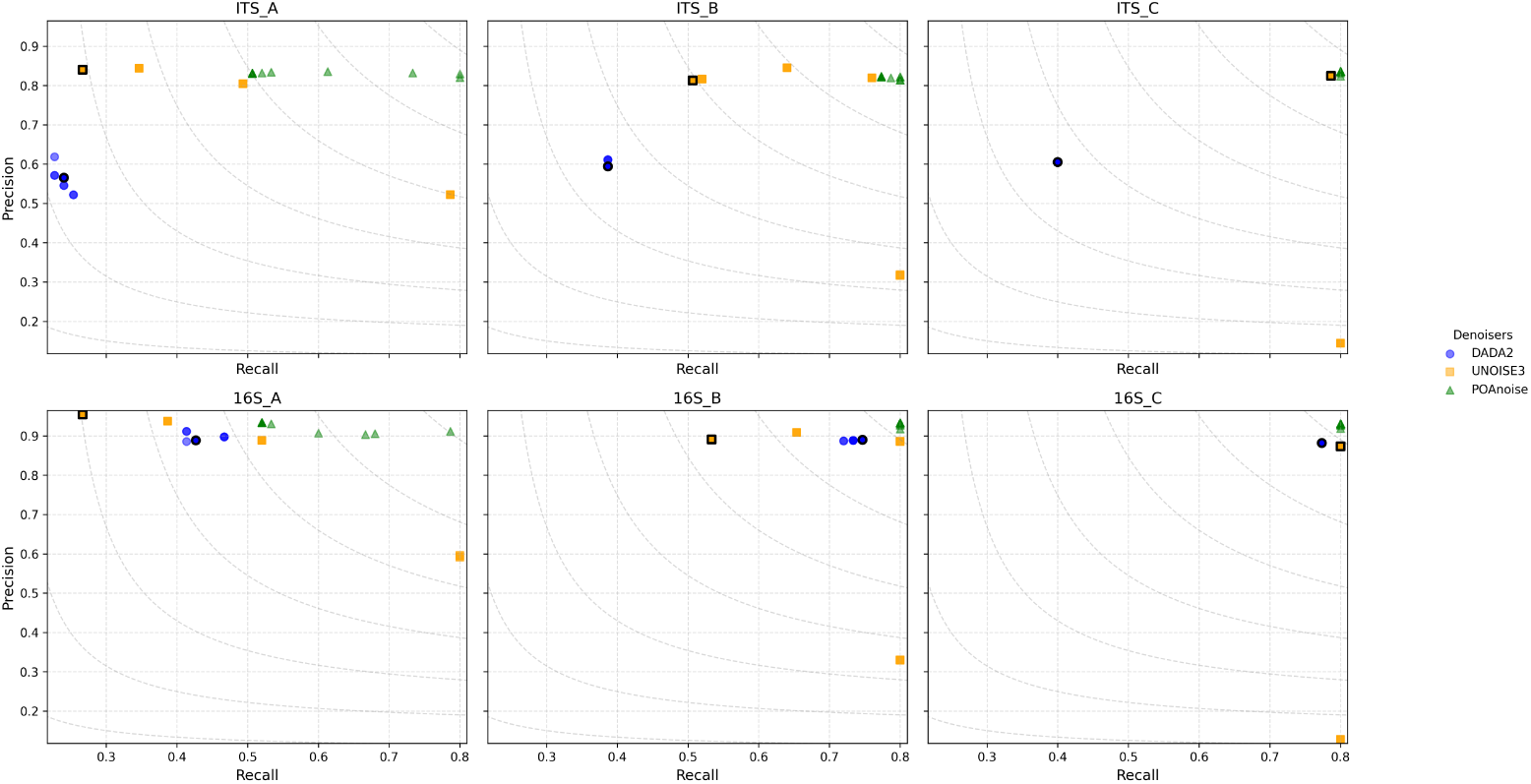
Precision–recall comparison of denoising methods across ITS and 16S datasets under varying parameter configurations. Each point represents one parameter configuration. Dashed gray lines indicate constant F_1_-score levels. The black outline identifies the recommended parameter configuration for DADA2 or UNOISE3.

The distributions of sequence error rates across parameter settings for the ITS and 16S datasets are shown in Figure 5. Across ITS datasets, both POAnoise and UNOISE3 consistently produced substantially lower error rates than DADA2. POAnoise generally showed comparable performance to UNOISE3, with slightly lower median error rates and narrower interquartile ranges in several datasets, indicating improved reconstruction stability. In contrast, DADA2 exhibited markedly elevated error rates across all ITS abundance regimes. For 16S datasets, the differences among denoisers were considerably smaller.

**Figure 5.**
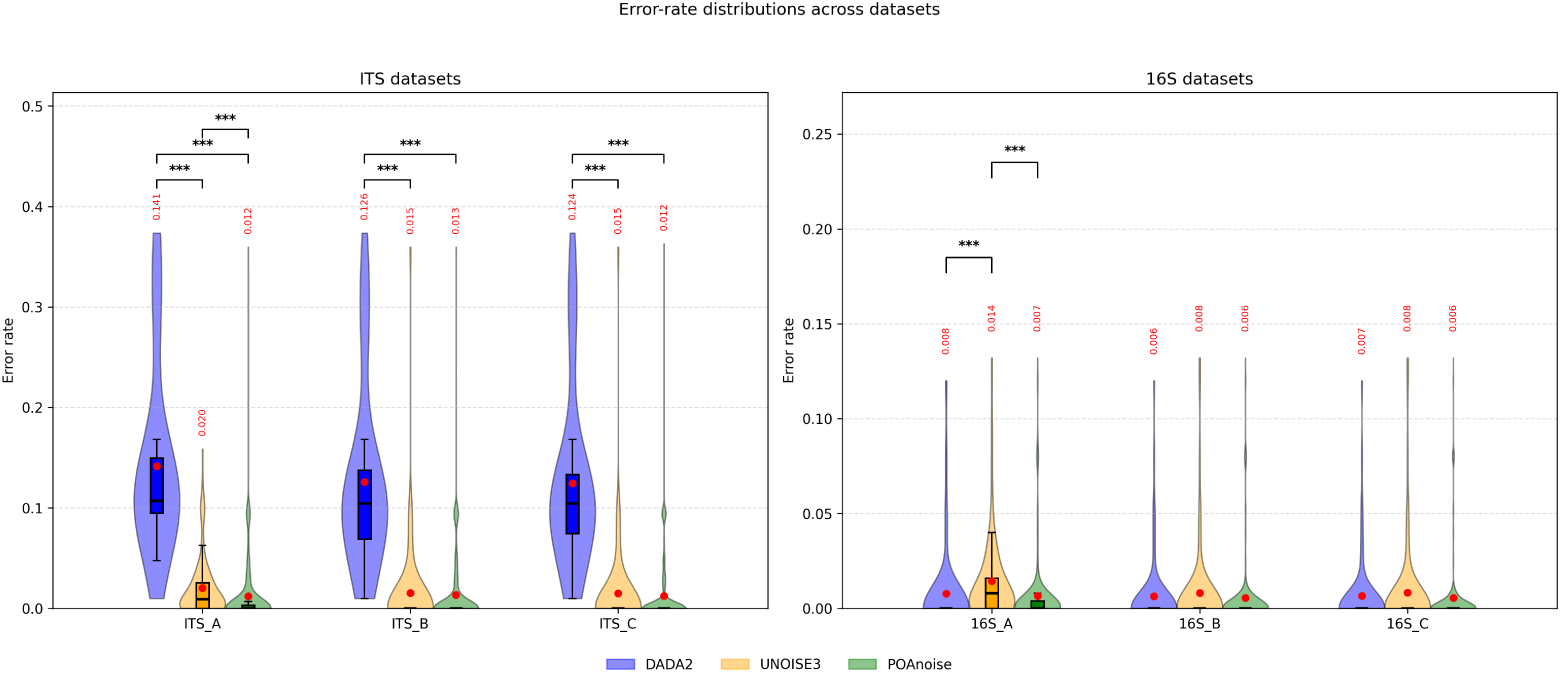
Distribution of sequence error rates across parameter settings for ITS (left) and 16S (right) datasets. Each violin shows the distribution of error rates across parameter configurations for a given denoising method and dataset. The embedded boxplot shows the median, IQR, and whiskers extending to the most extreme values within 1.5*×*IQR. Red dots denote the mean error rate, with the corresponding mean value shown in red above each distribution. Horizontal brackets indicate significant pairwise differences between denoising methods based on Holm-adjusted Mann–Whitney tests. One, two, and three stars indicate adjusted *p <* 0.05, *p <* 0.01, and *p <* 0.001, respectively; nonsignificant comparisons are not annotated.

Error-rate distributions differed substantially between marker types. ITS datasets showed broader error-rate distributions and larger upper whiskers, whereas 16S datasets exhibited more tightly concentrated low-error distributions. The greater sequence variability in ITS provides more sequence-level differentiation among the underlying reference sequences, which may facilitate distinguishing biological variants from sequencing errors. In contrast, the more conserved 16S sequences provide less sequence-level differentiation between variants.

Finally, abundance-aware evaluation revealed systematic differences in reconstruction across abundance classes (Figure 6). POAnoise maintained reconstruction ratios close to unity across abundances 4–256 (0.91–0.93), indicating relatively stable recovery of the expected abundance structure. UNOISE3 showed similar behavior, whereas DADA2 generally produced higher ratios (1.10–1.30). Because these ratios are calculated only for recovered reference sequences, ratios above one may reflect reads from unrecovered variants being attributed to recovered sequences, consistent with merging. The differences among denoisers were significant across abundance classes 4–256 (Kruskal–Wallis, *p <* 6 *×* 10*^−^*^5^), while the rarest class (*n* = 1) did not permit a comparable statistical analysis because of limited matched recovery.

**Figure 6.**
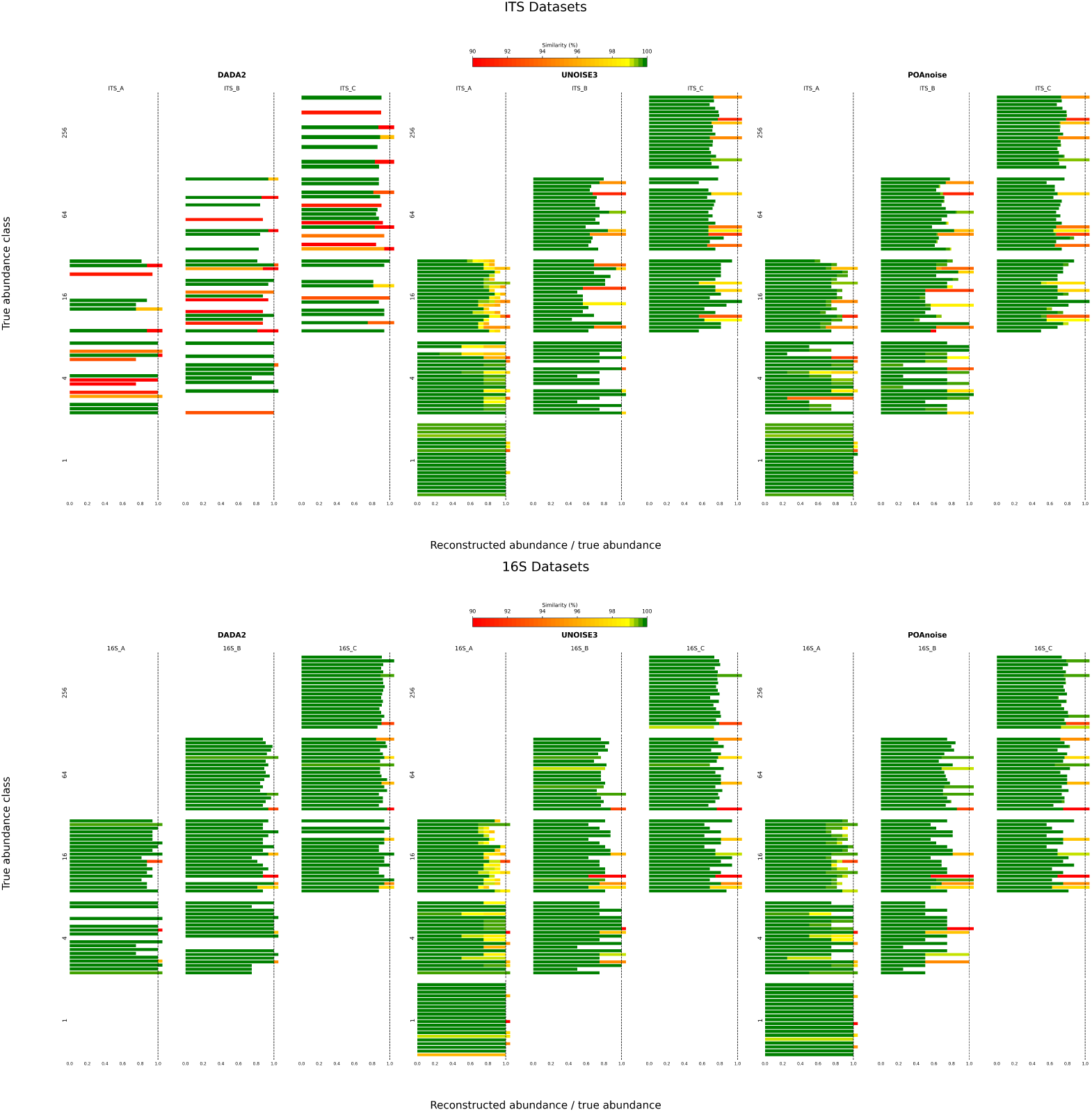
Abundance-aware denoising performance across ITS and 16S datasets. Rows correspond to true abundance classes (1, 4, 16, 64, or 256), while columns correspond to denoising methods. The horizontal axis shows the reconstructed abundance relative to the true abundance, with a value of 1 indicating exact abundance recovery. Ratios exceeding 1.05 are truncated at the plotting boundary and therefore may extend beyond the displayed range. Colors indicate the sequence similarity of the reconstructed sequences to their corresponding true reference sequences.

The computational cost of the three denoising methods differed substantially (Table 1). DADA2 and UNOISE3 completed the analyses within seconds to tens of seconds, whereas POAnoise required substantially longer runtimes, particularly as the number of input sequences increased. For the largest datasets, POAnoise required approximately 1.2–1.6 *×* 10^5^ s, compared with less than 41 s for DADA2 and 2.4 s for UNOISE3.

**Table 1.** Summary of dataset characteristics and denoising runtimes.

| # Sequences | # Species | Unique seq./species | DADA2<br>Runtime (s) | UNOISE3<br>Runtime (s) | POAnoise<br>Runtime (s) |
| --- | --- | --- | --- | --- | --- |
| 100 | 10 | 10 | 28.91 | 0.03 | 230.26 |
| 1000 | 10 | 100 | 31.17 | 0.11 | 3189.21 |
| 1000 | 100 | 10 | 5.57 | 0.07 | 3675.62 |
| 10000 | 10 | 1000 | 40.71 | 2.33 | 120341.93 |
| 10000 | 100 | 100 | 39.69 | 1.83 | 148497.45 |
| 10000 | 1000 | 10 | 33.66 | 1.71 | 158434.49 |

## 4 Discussion

This study presents POAnoise as a graph-based denoising framework that integrates partial-order alignment with consensus-based sequence reconstruction. Across the simulated ITS and 16S datasets considered here, POAnoise generally produced higher F_1_-scores and more stable performance across parameter configurations than DADA2 and UNOISE3. These results suggest that the POA-based representation can provide a useful alternative to existing denoising strategies.

A potential explanation for the observed performance is the type of sequence information used by POAnoise. DADA2 and UNOISE3 primarily distinguish candidate biological sequences from errors using pairwise sequence relationships together with probabilistic or abundance-based criteria, whereas POAnoise explicitly represents multiple sequences within a partial-order alignment graph. This representation captures shared alignment paths as well as branching patterns arising from sequence variation. Errors occurring independently across reads are therefore expected to produce less consistently supported branches than sequence differences that recur across multiple reads. The resulting graph structure provides an additional source of information for consensus reconstruction. However, the contribution of the POA representation itself cannot be separated from the other components of the pipeline in the present study and would require dedicated ablation experiments.

The results also showed differences between the ITS and 16S benchmarks. POAnoise maintained high performance across both marker types, while the relative differences among methods were generally smaller for the more conserved 16S datasets. The broader sequence variation observed in the ITS datasets provides greater sequence-level differentiation among reference sequences, which may provide more structural information for the alignment-based approach. These observations are consistent with the performance patterns observed in the benchmark, but further experiments using a wider range of biological sequence diversity would be needed to determine how broadly this behavior generalizes.

An important limitation of the current implementation is computational cost. The runtime analysis presented in the Results (Table 1) shows that POAnoise is substantially slower than both DADA2 and UNOISE3, particularly for larger datasets. Runtime also increased with the number of unique sequences per species, indicating that sequence heterogeneity and the resulting graph complexity contribute substantially to the computational burden. This computational cost is an important consideration for practical application of POAnoise to large amplicon datasets. The current implementation has not been extensively optimized, and several directions could improve scalability. These include parallelization of independent alignment operations, more efficient graph construction and data structures, graph pruning, and heuristic or accelerated sequence–graph alignment. Such improvements could reduce computational cost while retaining the alignment-based information used for reconstruction.

Overall, the results indicate that POAnoise can provide a useful alternative to existing denoising strategies, with higher and more consistent F_1_-scores across the simulated ITS and 16S datasets considered here. The main trade-off is substantially higher computational cost in the current implementation. Future work should therefore focus both on evaluating POAnoise on empirical datasets and on improving the computational efficiency of its graph-based alignment and consensus procedures.

## 5 Acknowledgments

The research was funded by the Research Council of Finland (grants no. 336212, 345110 and 21000068261) and the European Research Council (ERC) under the European Union’s Horizon 2020 research and innovation programme (grant agreement No 856506: ERC-synergy project LIFE-PLAN).

## 6 Author Contributions

Muhammad Ardiyansyah developed and implemented the method, performed the analysis, and wrote the manuscript. Brendan Furneaux proposed the research problem and contributed key methodological ideas. Otso Ovaskainen supervised the project and contributed to the study design and manuscript revision.

## Notes

### Competing Interest Statement

The authors have declared no competing interest.

## References

1. K. Abarenkov, A. Zirk, T. Piirmann, R. Pöhönen, F. Ivanov, R. H. Nilsson, and U. Kõljalg. UNITE general FASTA release for fungi 2. UNITE community, 2024.

2. A. Amir, D. McDonald, J. A. Navas-Molina, E. Kopylova, J. T. Morton, Z. Xu, R. Knight, and C. Lozupone. Deblur rapidly resolves single-nucleotide community sequence patterns. mSystems, 2(2), 2017.

3. M. Bandekar, K. D. More, S. C. Seleyi, N. Ramaiah, J. Kekäläinen, and J. Akkanen. Comparative analysis of microbiome inhabiting oxygenated and deoxygenated habitats using V3 and V6 metabarcoding of 16S rRNA gene. Marine Environmental Research, 199:106615, 2024.

4. B. J. Callahan, P. J. McMurdie, and S. P. Holmes. Exact sequence variants should replace operational taxonomic units in marker-gene data analysis. The ISME Journal, 11(12):2639–2643, 2017.

5. B. J. Callahan, P. J. McMurdie, M. J. Rosen, A. W. Han, A. J. Johnson, and S. P. Holmes. DADA2: High-resolution sample inference from illumina amplicon data. Nature Methods, 13(7):581–583, 2016.

6. P. J. Cock, C. J. Fields, N. Goto, M. L. Heuer, and S. E. Rice. The sanger fastq file format for sequences with quality scores, and the solexa/illumina fastq variants. Nucleic Acids Research, 38(6):1767– 1771, 2010.

7. F. Di Giallonardo, T. E. Schlub, M. Shi, and E. C. Holmes. Evaluating the accuracy of amplicon-based microbiome inference methods using simulated data. PLoS One, 14(5):e0215116, 2019.

8. R. C. Edgar. UPARSE: Highly accurate otu sequences from microbial amplicon reads. Nature methods, 10(10):996–998, 2013.

9. R. C. Edgar. UNOISE2: Improved error-correction for illumina 16s and its amplicon sequencing. bioRxiv, page 081257, 2016.

10. O. Gotoh. Optimal sequence alignment allowing for long gaps. Bulletin of Mathematical Biology, 52(3):359–373, 1990.

11. S. Holm. A simple sequentially rejective multiple test procedure. Scandinavian Journal of Statistics, 6(2):65–70, 1979.

12. W. H. Kruskal and W. A. Wallis. Use of ranks in one-criterion variance analysis. Journal of the American Statistical Association, 47(260):583–621, 1952.

13. C. Lee, C. Grasso, and M. F. Sharlow. Multiple sequence alignment using partial order graphs. Bioinformatics, 18(3):452–464, 2002.

14. H. B. Mann and D. R. Whitney. On a test of whether one of two random variables is stochastically larger than the other. The Annals of Mathematical Statistics, 18(1):50–60, 1947.

15. A. E. Minoche, J. C. Dohm, and H. Himmelbauer. Evaluation of genomic high-throughput sequencing data generated on illumina HiSeq and genome analyzer systems. Genome biology, 12(11):R112, 2011.

16. S. B. Needleman and C. D. Wunsch. A general method applicable to the search for similarities in the amino acid sequence of two proteins. Journal of Molecular Biology, 48(3):443–453, 1970.

17. C. Quast, E. Pruesse, P. Yilmaz, J. Gerken, T. Schweer, P. Yarza, J. Peplies, and F. O. Glöckner. The SILVA ribosomal RNA gene database project: improved data processing and web-based tools. Nucleic Acids Research, 41(D1):D590–D596, 2013.

18. F. J. Rang, W. P. Kloosterman, and J. de Ridder. From squiggle to basepair: computational approaches for improving nanopore sequencing read accuracy. Genome Biology, 19(1):90, 2018.

19. M. J. Rosen, B. J. Callahan, D. S. Fisher, and S. P. Holmes. Denoising PCR-amplified metagenome data. BMC Bioinformatics, 13(1):283, 2012.

20. P. D. Schloss, S. L. Westcott, T. Ryabin, J. R. Hall, M. Hartmann, E. B. Hollister, R. A. Lesniewski, B. B. Oakley, D. H. Parks, C. J. Robinson, et al. Introducing mothur: open-source, platform-independent, community-supported software for describing and comparing microbial communities. Applied and environmental microbiology, 75(23):7537–7541, 2009.

21. S. S. Shapiro and M. B. Wilk. An analysis of variance test for normality (complete samples). Biometrika, 52(3–4):591–611, 1965.

22. R. Vaser, I. Sović, N. Nagarajan, and M. Sikić. Fast and accurate de novo genome assembly from long uncorrected reads. Genome Research, 27(5):737–746, 2017.

23. P. Yilmaz, L. W. Parfrey, P. Yarza, J. Gerken, E. Pruesse, C. Quast, T. Schweer, J. Peplies, W. Ludwig, and F. O. Glöckner. The SILVA and ”All-species Living Tree Project (LTP)” taxonomic frameworks. Nucleic Acids Research, 42(D1):D643–D648, 2014.

